# Allosteric Inhibition of CRISPR-Cas9 by Bacteriophage-derived Peptides

**DOI:** 10.1101/642538

**Authors:** Yan-ru Cui, Shao-jie Wang, Jun Chen, Jie Li, Wenzhang Chen, Shuyue Wang, Bing Meng, Wei Zhu, Zhuhong Zhang, Bei Yang, Biao Jiang, Guang Yang, Peixiang Ma, Jia Liu

## Abstract

**Background:** CRISPR-Cas9 has been developed as a therapeutic agent for various infectious and genetic diseases. In many clinically relevant applications, constitutively active CRISPR-Cas9 is delivered into human cells without a temporal control system. Excessive and prolonged expression CRISPR-Cas9 can lead to elevated off-target cleavage. The need for modulating CRISPR-Cas9 activity over the dimensions of time and dose has created the demand of developing CRISPR-Cas off-switches. Protein and small molecule-based CRISPR-Cas inhibitors have been reported in previous studies.

**Results:** We report the discovery of Cas9-inhibiting peptides from inoviridae bacteriophages. These peptides, derived from the periplasmic domain of phage major coat protein G8P (G8P_PD_), can inhibit the *in vitro* activity of *Streptococcus pyogenes* Cas9 (SpCas9) proteins in an allosteric manner. Importantly, the inhibitory activity of G8P_PD_ on SpCas9 is dependent on the order of guide RNA addition. Ectopic expression of full-length G8P (G8P_FL_) or G8P_PD_ in human cells can inactivate the genome-editing activity of SpCas9 with minimum alterations of the mutation patterns. Furthermore, unlike the anti-CRISPR protein AcrII4A that completely abolishes the cellular activity of CRISPR-Cas9, G8P co-transfection can reduce the off-target activity of co-transfected SpCas9 while retaining its on-target activity.

**Conclusion:** G8Ps discovered in the current study represent the first anti-CRISPR peptides that can allosterically inactivate CRISPR-Cas9. This finding may provide insights into developing next-generation CRISPR-Cas inhibitors for precision genome engineering.

## Background

Clustered Regularly-Interspaced Short Palindromic Repeats (CRISPR) is the bacterial adaptive immune system to defend bacteriophage infections [1–3]. During infection, invader DNA is captured and integrated into bacterial genome as CRISPR array. Sequences from CRISPR array are transcribed and processed into CRISPR RNAs (crRNAs), which direct CRISPR-associated (Cas) proteins to foreign nucleic acids [1, 2]. Type II CRISPR-Cas systems function with streamlined components comprising of a single nuclease protein such as Cas9 [4]. The modular and programmable features make CRISPR-Cas9 one of the most widely used tools for genome engineering applications [5–8]. However, CRISPR-Cas9 is associated with off-target cleavage [9], chromosomal rearrangement [10] and genotoxicity [11]. These side effects mainly arise from the excessive or prolonged expression of CRISPR-Cas9 [12–14]. As a therapeutic agent, CRISPR-Cas9 is often constitutively expressed in host cells [15], making the elevated off-activity a major safety concern. Temporal control of SpCas9 activity have been investigated as an approach to improving its specificity in human cells. Technologies enabling the temporal control of CRISPR-Cas9 include optogenetic tools, intein splicing system, small molecule inducers or inhibitors [16–20] and anti-CRISPR proteins (Acrs) [21, 22].

The most investigated CRISPR-Cas inhibitors are the naturally occurring, phage-derived Acrs. Bacteriophages can use Acrs to antagonize the CRISPR-Cas immunity in bacteria [23, 24]. A number of Acrs have been identified for type I [23, 25–28], type II [29–32] and type V [28, 33] CRISPR-Cas systems. Acrs can be adapted to regulate CRISPR-Cas activities in bacteria [34], yeast [35] and mammalian cells [29, 31, 34, 36–38]. Biosensor [39] and synthetic circuits [40] can be devised based on Acr-coupled CRISPR-Cas systems. Moreover, Acrs can be harnessed to enable the temperature-responsive [41] and optogenetic [42] control of CRISPR-Cas activity. Importantly, Acrs can enhance the editing [38] and cell-type [43] specificities and reduce the cytotoxicity [11] of CRISPR-Cas-mediated genome editing. It has also been reported that Acrs can facilitate the production of CRISPR-carrying viral vectors by restricting CRISPR self-cleavage [44].

Currently known Acrs inhibit CRISPR-Cas systems by interfering with Cas protein-mediated DNA surveillance or cleavage [21]. For instance, AcrIIA4 mimics double-stranded DNA (dsDNA) and occupies the protospacer adjacent motif (PAM) recognition site of SpCas9, thereby preventing Cas9 protein from binding to the target DNA [38, 45, 46]. Using a different mechanism, AcrIIC3 perturbs DNA binding by inducing the dimerization of Cas9 protein [36]. An alternative Cas inactivation strategy by Acrs is to interact with DNA-bound Cas proteins and block subsequent DNA cleavage, as seen with AcrF3 [47, 48] and AcrIIC1 [36]. In addition, some Acrs can function as acetyltransferase and inactivate CRISPR-Cas activity by post-translational modifications [49]. Different Acrs may inactivate CRISPR-Cas via identical mechanisms while possessing low sequence similarities [21]. The Acrs characterized to date share no common sequence motifs except for a putative transcriptional element referred to as anti-CRISPR associated genes (Acas) that are commonly found downstream of the Acr genes in the bacteriophage genome [26]. The poorly understood sequence-structure-activity relationship largely hampers the systemic discovery of novel Acrs. In addition to protein-based inhibitors, small-molecule CRISPR-Cas inhibitors have been developed using a high-throughput screening platform [20]. These small molecules are cell-permeable and can reversely disrupt SpCas9-DNA interaction, thus enabling dose and temporal control of SpCas9. However, relatively high concentrations of 10 μM or above are required for small molecules to achieve efficient inhibition [20].

Along with small molecules and proteins, peptides represent an alternative class of CRISPR-inhibiting agents with distinct biochemical features. In this study, we report the discovery of Cas9-inactivating peptides from inoviridae bacteriophages. In an attempt to develop anti-CRISPR antibodies using well established phage display technology [50], we are surprised to find that the commonly used laboratory bacteriophage strain M13 served as a source of Cas9-inactivating agent. Subsequent analyses showed that the periplasmic domain of the major coat protein G8P (G8P_PD_) from several inoviridae bacteriophages, which contain M13 phage, inhibited the *in vitro* and *in vivo* activity of SpCas9 in an allosteric manner. Our study hence expands the inhibitor toolbox for the temporal control of CRISPR-Cas activity.

## Results

### Intact M13 phage inhibits the *in vitro* DNA cleavage activity of SpCas9

In a conventional phage display experiment, we surprisingly discovered that intact M13 phage [51] itself could block the DNA cleavage activity of purified SpCas9 proteins in dose-dependent manner with an approximate half maximum inhibitory concentration (IC_50_) of 5 nM (Fig. 1a), which corresponds to a phage titer of 3×10^9^ PFU/μL. Interestingly, phage-mediated SpCas9 inactivation occurred only if phage was supplemented to the reaction prior to the addition of sgRNA, but not post the formation of SpCas9-sgRNA ribonucleoproteins (RNPs) (Fig. 1b). The order-of-addition-dependent inhibition suggested that competition for sgRNA-binding site in SpCas9 is a possible mechanism of the inhibitory activity of M13 phage.

**Fig. 1.**
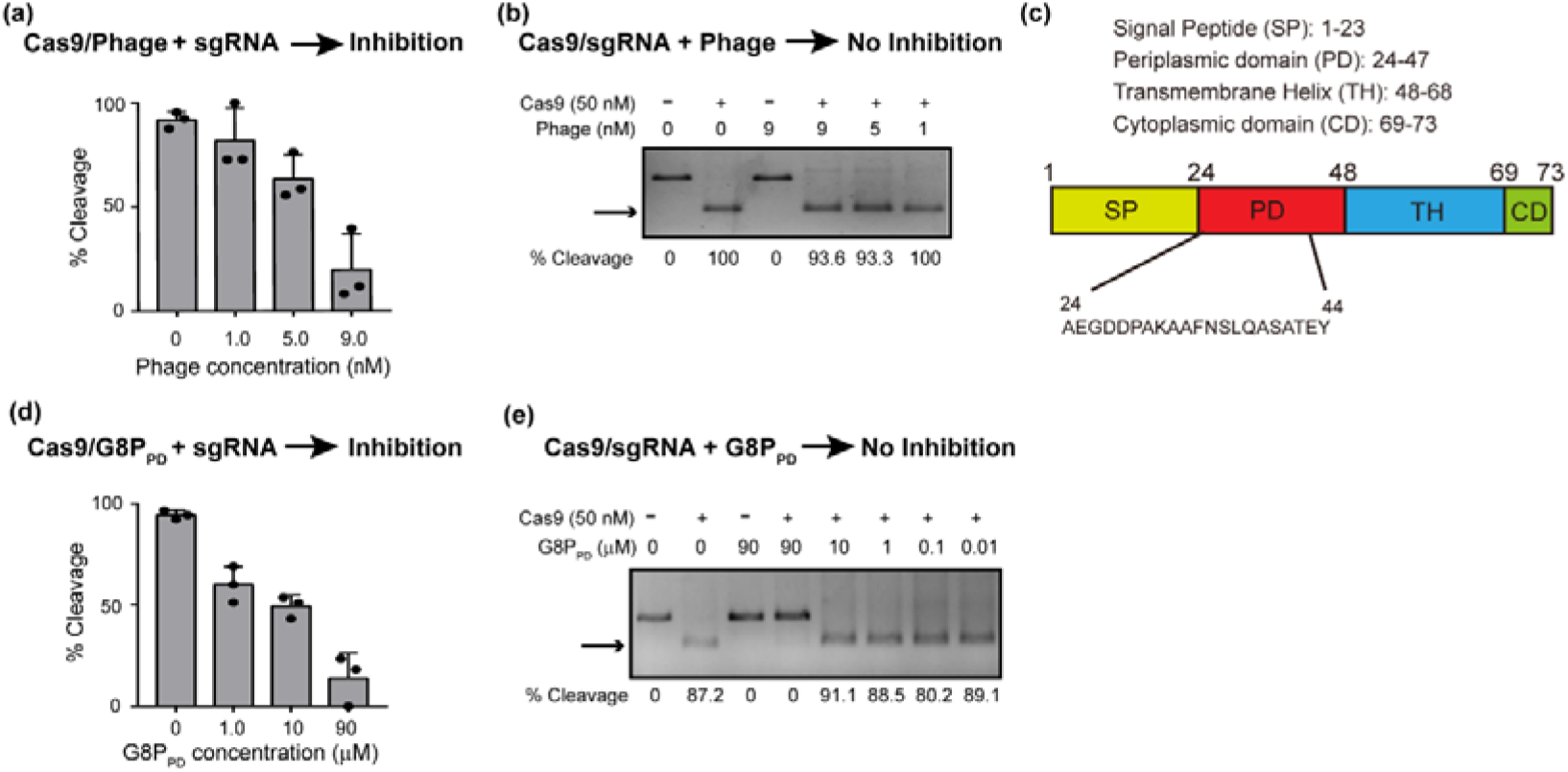
Inhibition of the *in vitro* activity of SpCas9 by intact M13 phage and phage-derived G8P_PD_ peptides. (**a**) Dose-dependent inhibition of SpCas9 by intact M13 phage. (**b**) Intact M13 phage does not inhibit the *in vitro* activity of assembled Cas9-sgRNA RNP. (**c**) Structural organization of M13 phage major coat protein G8P. (**d**) Dose-dependent inhibition of SpCas9 by G8P_PD_. (**e**) G8P_PD_ does not inhibit the *in vitro* activity of assembled Cas9-sgRNA RNP. The above reactions are performed in the absence or presence of 50 nM SpCas9 proteins. The results are shown as mean ± SD (*n* = 3). Arrows indicate cleavage products.

### M13 phage major coat protein-derived peptide inhibits the *in vitro* DNA cleavage activity of SpCas9

Next we sought to determine the components in M13 phage that contribute to SpCas9 inactivation. M13 phage has a simple, compact genome comprising of 11 protein-coding sequences. Considering the accessibility, surface-exposed phage proteins most likely serve as the sources of Cas9-inhibiting agents. Fully packaged M13 phage contains approximately 2700 copies of major coat protein pVIII (G8P) and 5 copies each of minor coat proteins pIII, pVI, pVII and pXI [52–54]. We initiated our investigation on the major coat protein G8P due to its abundance on M13 phage surface. The 73-amino acid major coat protein G8P contains four segments including signal peptide, periplasmic domain, transmembrane helix and cytoplasmic domain (Fig. 1c). After G8P maturation, signal peptide is cleaved and transmembrane helix is inserted into phage membrane, leaving periplasmic domain the only region on phage surface. Therefore, we synthesized a 21-amino acid peptide constituting the periplasmic domain of G8P (G8P_PD_) and examined its inhibitory activity on SpCas9.

We found that G8P_PD_ inhibited the activity of SpCas9 with an IC_50_ of 5 μM (Fig. 1d), which was 1000-fold lower than that of intact M13 phage. Similar to the intact phage, G8P_PD_ suppressed the *in vitro* DNA cleavage of SpCas9 in an order-of-addition-dependent manner (Fig. 1e), indicating that G8P_PD_ may inactivate SpCas9 by specifically interfering with apo-SpCas9.

To explore whether peptides with sequence similarity to G8P_PD_ can inhibit Cas9 activity, we used Blast to search for peptide homologs of M13 G8P_PD_. We identified several peptide sequences from inoviridae phages (see below for detailed characterization). Interestingly, G8P_PD_ peptides from f1 and M13 phages have only one amino acid difference (Additional file 1: Figure S1a) and similar activities on inhibiting Cas9 cleavage or Cas9-sgRNA binding (Additional file 1: Figure S1b-c). The synthetic f1 G8P_PD_ oligopeptide seemed to have higher aqueous solubility and is used interchangeably with M13 G8P_PD_ in the following studies.

### G8P_PD_ prevents the assembly of Cas9 and sgRNA by binding to sgRNA-free Cas9 (apo-Cas9)

SpCas9 can be inactivated at distinct steps during its action including guide RNA binding, substrate DNA binding and DNA cleavage. Most previously known Acrs exert their inhibitory activity by interfering with DNA surveillance or cleavage. To understand the mechanism of actions of G8P_PD_, we examined the interactions between G8P_PD_ and apo-Cas9 or Cas9-sgRNA RNP using electrophoresis mobility shift assay (EMSA). Under fixed sgRNA concentration of 15 μM, the fraction of Cas9-bound sgRNA increased with the increasing molar ratio of Cas9 and sgRNA in the absence of G8P_PD_. By contrast, pre-incubation of SpCas9 proteins with 300 μM G8P_PD_ reduced the fraction of Cas9-bound sgRNA in a G8P_PD_ dose-dependent manner (Fig. 2a). We also noticed that under high molar ratio of Cas9 and sgRNA, G8P_PD_ did not fully block the formation of Cas9-sgRNA RNP (Fig. 2a), suggesting a weak interaction between Cas9 and G8P_PD_ that is reversible under high concentrations of sgRNA.

**Fig. 2.**
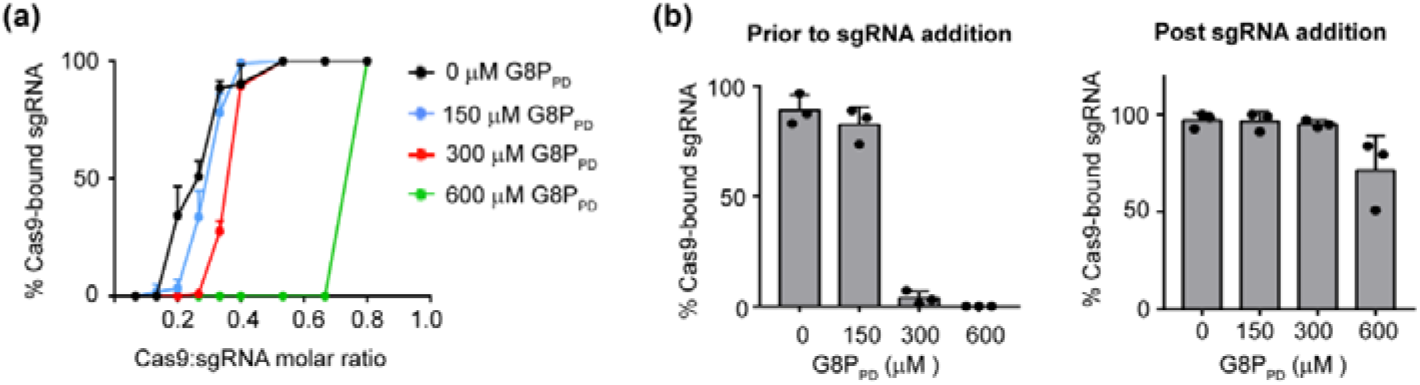
G8P_PD_ prevents Cas9-sgRNA assembly. (**a**) Dose-dependent inhibition of Cas9-sgRNA binding by f1 G8P_PD_. sgRNA concentration is fixed to 15 μM. (**b**) f1 G8P_PD_ prevents Cas9-sgRNA assembly prior to, but not post, sgRNA addition. Cas9 to sgRNA ratio is fixed to 0.3. The above results are shown as mean ± SD (*n* = 3).

G8P_PD_-mediated inhibition of Cas9-sgRNA assembly is dependent on the order of sgRNA addition. When the ratio of the molar concentrations of Cas9 and sgRNA is fixed to 1: 0.3, pre-incubation of Cas9 with 300 or 600 μM f1 G8P_PD_ prior to sgRNA addition abolished the assembly of Cas9 and sgRNA (Fig. 2b). By contrast, supplementation of 300 or 600 μM G8P_PD_ post sgRNA addition had minor or no effect on the formation of Cas9-sgRNA complex (Fig. 2b). These results suggest that G8P_PD_ prevents Cas9-sgRNA binding by interacting with apo-Cas9 but not sgRNA-bound Cas9 and may explain why G8P_PD_-mediated inactivation of SpCas9 cleavage is dependent on the order of sgRNA addition.

### Identification of G8P_PD_ binding site in SpCas9

In order to dissect the mechanism of interactions between SpCas9 and G8P_PD_, we sought to determine the binding region of G8P_PD_ on SpCas9 using high-resolution mass spectrometry (MS). SpCas9 protein and M13 G8P_PD_ were crosslinked using collision-induced dissociation (CID)-cleavable cross-linker disuccinimido sulfoxide (DSSO) [55] and then were subject to digestion with chymotrypsin. The integration analyses of CID-induced cleavage of interlinked peptides in MS/MS and MS^3^ of single peptide chain fragment ions revealed high crosslinking scores (Fig. 3a) on residues K1158 of [K]SVKEL peptide and K1176 of E[K]NPIDFLEAKGY peptide from SpCas9 (Fig. 3b-c and Additional file 1: Figure S2a-b). These peptides occupy a continuous region in the PAM-interacting (PI) domain of SpCas9 (Fig. 3d) that is responsible for recognizing the PAM sequence on the non-complementary DNA strand [56]. Interestingly, this candidate G8P_PD_ binding site dose not locate in the sgRNA or DNA binding pockets of SpCas9. These results suggest that G8P_PD_ does not directly compete with sgRNA but may instead function as an allosteric inhibitor.

**Fig. 3.**
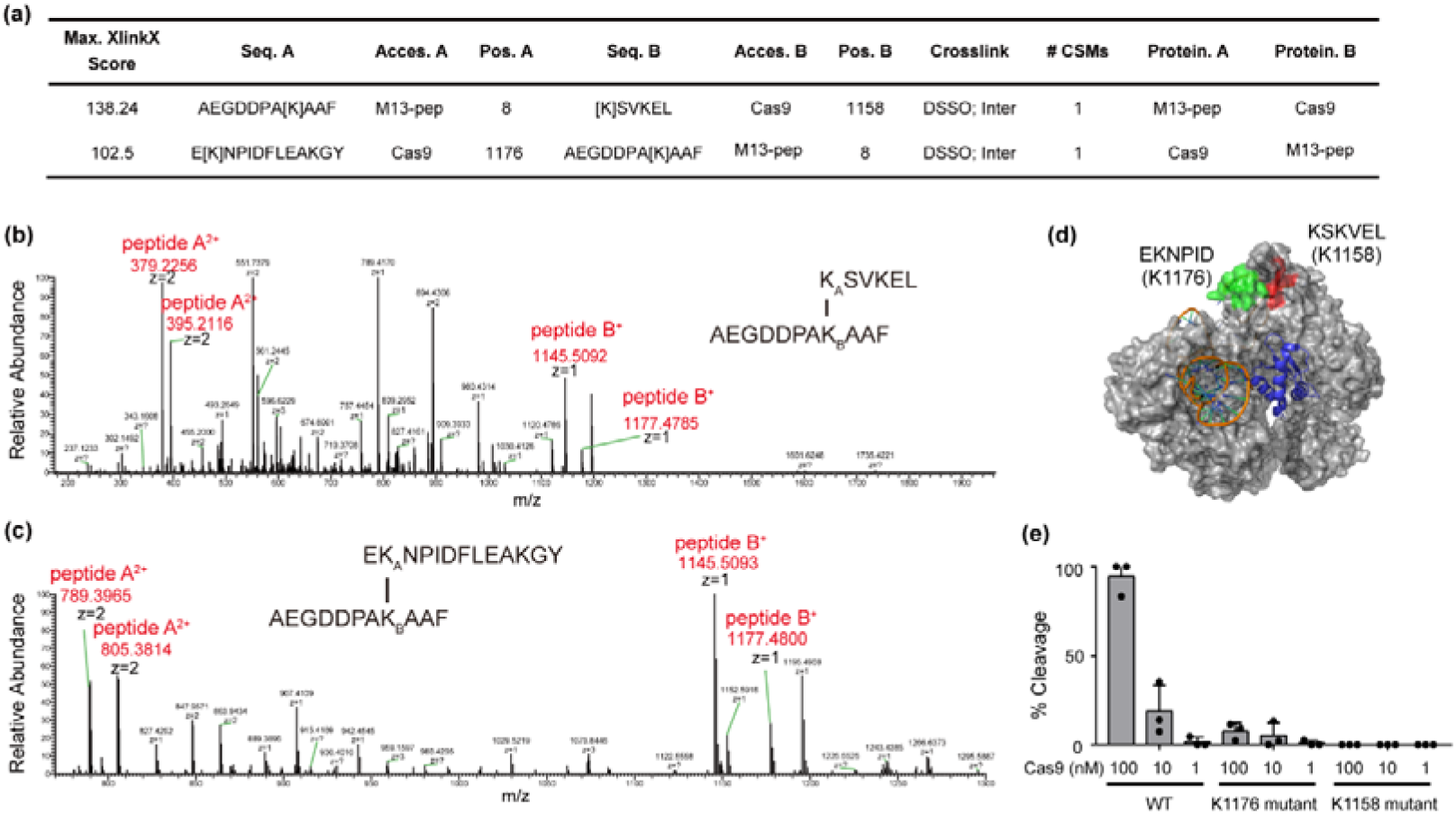
Identification of M13 G8P_PD_ binding site in the PI domain of SpCas9. (**a**) Maximum XlinkX scores of peptide hits in high-resolution MS analyses. (**b**) Secondary MS showing crosslinked peptides KSVKEL-AEGDDPAKAAF. (**c**) Secondary MS showing crosslinked peptides EKNPIDFLEAKGY-AEGDDPAKAAF. (**d**) Location of G8P_PD_ binding sites in SpCas9. The structure of SpCas9 in complex with AcrIIA4 (5VW1) is displayed by PyMOL. AcrIIA4 is shown in blue. The candidate G8P_PD_ binding sites on SpCas9 are shown in green and red respectively. (**e**) *In vitro* DNA cleavage by WT, K1176 mutant and K1158 mutant SpCas9. Arrow indicates cleavage product. The results are shown as mean ± SD (*n* = 3).

Next we sought to perform mutational analyses on the candidate G8P_PD_ binding sites in SpCas9. Residues KSKVEL in K1158 mutant and EKNPID in K1176 mutant were mutated into alanines respectively. Mutant SpCas9 proteins were purified into high homogeneity (Additional file 1: Figure S3a-b). *In vitro* cleavage reaction illustrated that alanine mutations at positions KSKVEL (K1158 mutant) and EKNPID (K1176 mutant) markedly reduced the DNA cleavage activity of SpCas9 (Fig. 3e), suggesting the importance of the G8P_PD_ binding sites for SpCas9 activity.

### **α**-helical structure is critical for the inhibitory activity of G8P_PD_

Next we performed an alanine scan on f1 phage G8P_PD_ to illustrate its structural determinants for Cas9 inhibition. Four peptide mutants are designed with alanine mutations spanning the entire G8P_PD_ sequence (Fig. 4a). Alanine mutations at positions 6 to 11 abolished the inhibitory activity of G8P_PD_ while mutants 1, 3 and 4 retained the majority of Cas9-inhibiting activity (Fig. 4a). The major coat protein G8P adopts an α-helical structure [54]. Residues 6 to 11 are located at the N-terminus of the α helix and contain Pro6, Lys8, Phe11 and three native alanines (Fig. 4b). The abolished activity in mutant 2 suggested a critical role of positions 6 to 11, particularly the residues Pro6, Lys8 and Phe11. Circular dichroism (CD) spectra study revealed an α-helical structure-enriched feature for f1 G8P_PD_ WT but not for mutant 2 (Fig. 4c), suggesting that Pro6, Lys8 and Phe11 are important for maintaining the α-helical structure in f1 G8P_PD_. Although the presence of residues 6 to 11 could directly participate in the interaction between Cas9 and G8P_PD_, the overall α-helical structure of G8P_PD_ could be also important for its Cas9-inhibiting activity.

**Fig. 4.**
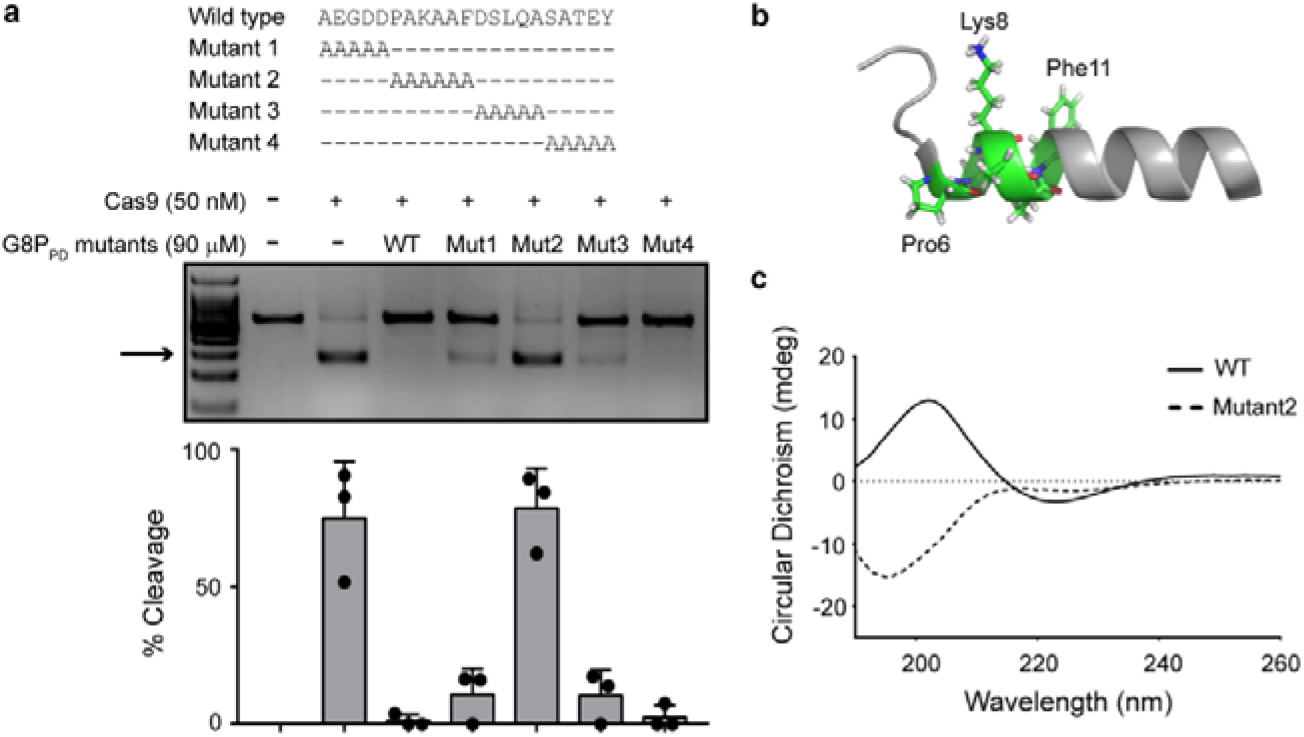
α-helical structure is critical for the Cas9-inhibiting activity of G8P_PD_. (**a**) Alanine mutations at position 6-11 abolish the inhibitory activity of G8P_PD_. The results are shown as mean ± SD (*n* = 3). The arrow denotes cleavage products. (**b**) Structure of G8P_PD_ peptide (PDB entry 2MJZ), displayed by PyMOL. Residues 6-11 are shown as stick. (**c**) CD spectra of f1 G8P_PD_ WT and mutant 2.

### Modulation of SpCas9 activity in human cells using inoviridae phage G8P_PD_

We next explored the potential application of G8P_PD_ as an off-switch for the genome-editing activity of SpCas9 in human cells. Ectopic expression of the full-length (G8P_FL_) or periplasmic domain (G8P_PD_) of M13 and f1 G8Ps at 24 h prior to Cas9-sgRNA transfection significantly suppressed the genome-editing activity of SpCas9 in HEK293 cells (Fig. 5a-b). In comparison, Acr protein AcrII4A [38] blocked SpCas9 cleavage on *AAVS1* site whereas *Neisseria meningitidis* Cas9 (NmeCas9)-specific Acr protein AcrIIC3 [57] partially inhibited SpCas9. Importantly, f1 G8P_PD_ was capable to inhibit SpCas9 activity across different genes and cell types (Fig. 5c). Consistent with the *in vitro* experiments, significant inhibition of the on-target activity of SpCas9 in human cells was observed only when G8P_PD_ was overexpressed prior to sgRNA transfection. Co-transfection of G8P_PD_ and SpCas9-sgRNA did not inhibit SpCas9 cleavage (*P*>0.05) (Fig. 5d).

**Fig. 5.**
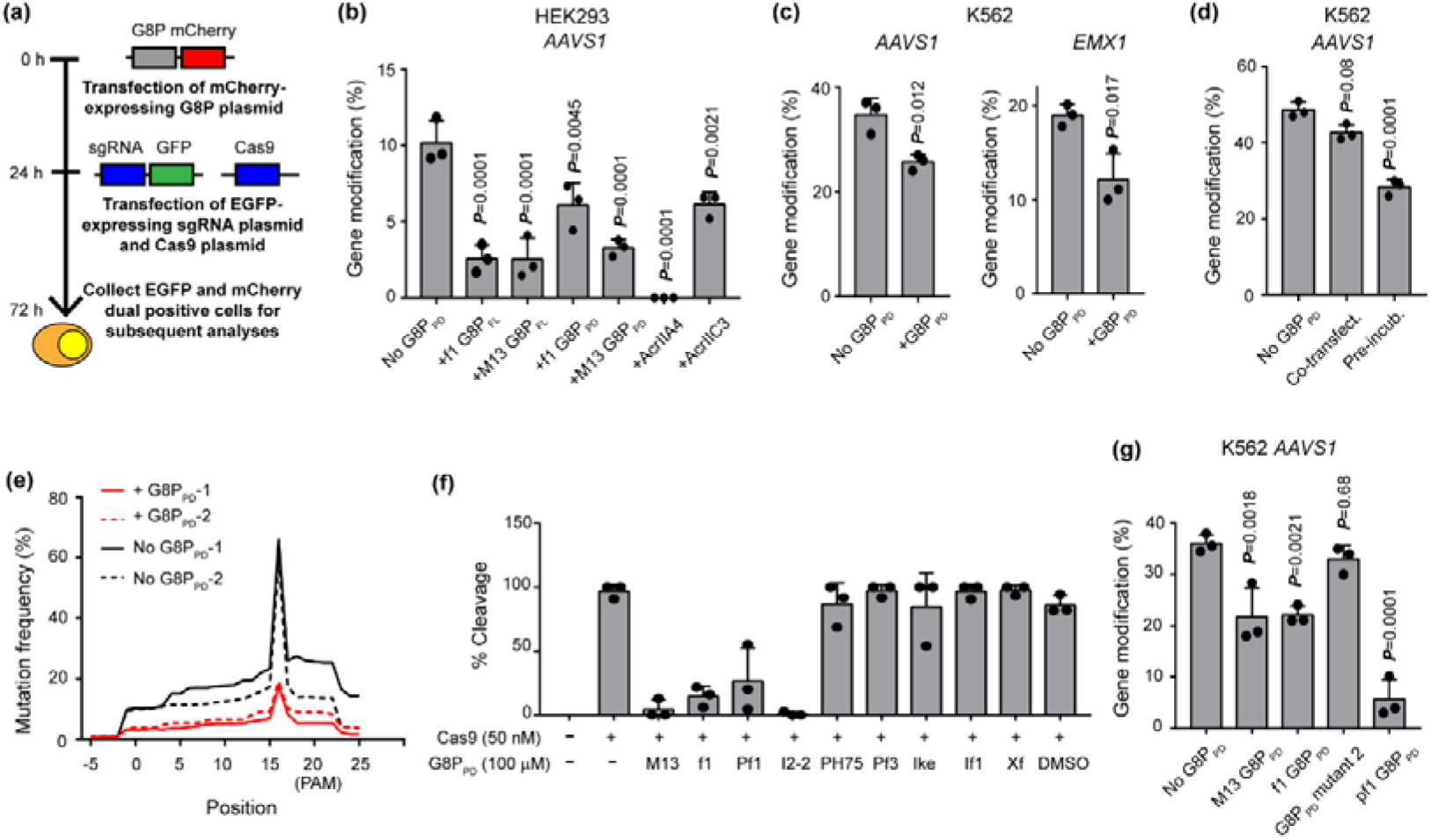
Inhibition of the genome-editing activity of SpCas9 in human Cells by inoviridae phage G8P_PD_. (a) Flowchart showing experimental design. (**b**) Comparison of SpCas9-inhibiting activities of phage peptides and Acrs in HEK293 cells. (**c**) G8P_PD_ inhibits the genome-editing activity of SpCas9 across different genes and cell types. (**d**) The effects of G8P_PD_ co-transfection and pre-incubation on the genome-editing activity of CRISPR-Cas9. (**e**) Density plot showing the NGS analyses of the distribution of mutation rates along the edited genomic sites of *AAVS1* in K562 cells. The results of two biological replicates are individually shown. (**f**) Inhibition of the *in vitro* DNA cleavage activity of SpCas9 by inoviridae G8P_PD_. DMSO of 0.1% is included as a solvent control. (**g**) Inhibition of SpCas9 activity in K562 cells by inoviridae G8P_PD_. The results are shown as mean ± SD (*n* = 3). Significant difference between test groups and mock is determined by one-way ANOVA with Dunnett’s multiple comparisons test. The adjusted *P* values are indicated.

In order to have detailed understanding of the effects of G8P_PD_ on the genome-editing activity of SpCas9, we performed next-generation sequencing (NGS) to analyze the profiles of edited genomic loci in the absence and presence of G8P_PD_. Despite reduced mutation rate, the mutation pattern of SpCas9 along the 20 bp sgRNA-targeting site was not altered by G8P_PD_ treatment, as characterized by the high-frequency editing events at 3 bp upstream of the PAM sequence [4] (Fig. 5e). Importantly, G8P_PD_ treatment retained the distribution pattern of indel length, with 1-5 bp indel being predominant in the population (Additional file 1: Figure S4a). In addition, we observed modest decrease in the in-frame mutations (3N) (Additional file: Figure S4b), the mechanism of which is yet to be elucidated. Collectively, these data suggested that G8P_PD_ treatment did not cause major alterations in the profiles of SpCas9-induced mutations, thus highlighting the potential of G8P_PD_ as a safe off-switch for the therapeutic applications of SpCas9.

To expand peptide-based anti-CRISPR toolbox, we examined the G8Ps from other inoviridae phages (Additional file 1: Figure S5). Peptides constituting the periplasmic domain of these G8P (G8P_PD_) are synthesized and evaluated for the *in vitro* and *in vivo* activities. At a concentration of 100 μM, the G8P_PD_ from M13, f1, Pf1 and I2-2 phage markedly inhibited the *in vitro* DNA cleavage activity of SpCas9 while other G8P_PD_ orthologues showed little inhibitory effects (Fig. 5f). Ectopic expression of M13, f1 and pf1 G8P_PD_ in K562 cells significantly reduced the activity of SpCas9 in HEK293 cells whereas G8P_PD_ mutant 2 did not show inhibitory activity (*P*>0.05) (Fig. 5g). Our results suggested that inoviridae phage G8Ps could be leveraged to inhibit both the *in vitro* and *in vivo* activity of SpCas9. The variations of G8P_PD_ sequences and the difference in their inhibitory activities indicate that further engineering endeavors could be made to improve Cas9-inhibiting peptides.

To understand the specificity of G8P_PD_ within type II CRISPR system, we examine the inhibitory activity of f1 G8P_PD_ on Cas9 orthologues from *Staphylococcus aureus* (SaCas9) and *Neisseria meningitidis* (NmCas9). NGS analyses showed that pre-incubation of HEK293 cells with G8P_PD_ did not significantly inhibit the activities of NmCas9 or SaCas9 at the examined genomic sites (Additional file: Figure S6).

### G8P co-transfection improves the specificity of SpCas9 in human cells

It has been reported that timed delivery of AcrIIA4 can improve the genome-editing specificity of nucleofected Cas9-sgRNA RNP complex [38]. We intended to investigate the effects of G8P peptides on the specificity of constitutively expressed SpCas9, a more therapeutically relevant model. The aforementioned *in vitro* and *in vivo* data have suggested that the inhibitory effects of G8P on CRISPR-Cas9 is dependent on the accessibility of Cas9 protein to sgRNA. This observation prompted us to explore whether G8P can be leveraged, via timed delivery, to improve the specificity of SpCas9. Unlike the above experiments using G8Ps as CRISPR-Cas9 off switch (Fig. 5), here we co-transfected Hela cells with Acr or G8P plasmid and sgRNA and SpCas9-coding plasmids. AcrIIA4 suppressed the on- and off-activities of SpCas9 to undetectable levels, as determined by T7E1 assay (Fig. 6a). NGS analyses showed that M13 G8P_FL_ and f1 G8P_FL_ significantly reduced (*P*<0.05) both on-target and off-target activity of SpCas9 in Hela cells. It appeared that the inhibitory effects are more prominent at the off-target site (Fig. 6b). AcrIIC3 significantly reduced the on-target activity (*P*<0.05) but not the off-target activity (*P*>0.05) (Fig. 6b). Importantly, M13 G8P_PD_ could reduce the off-target events without affecting the on-target cleavage (Fig. 6a-b).

**Fig. 6.**
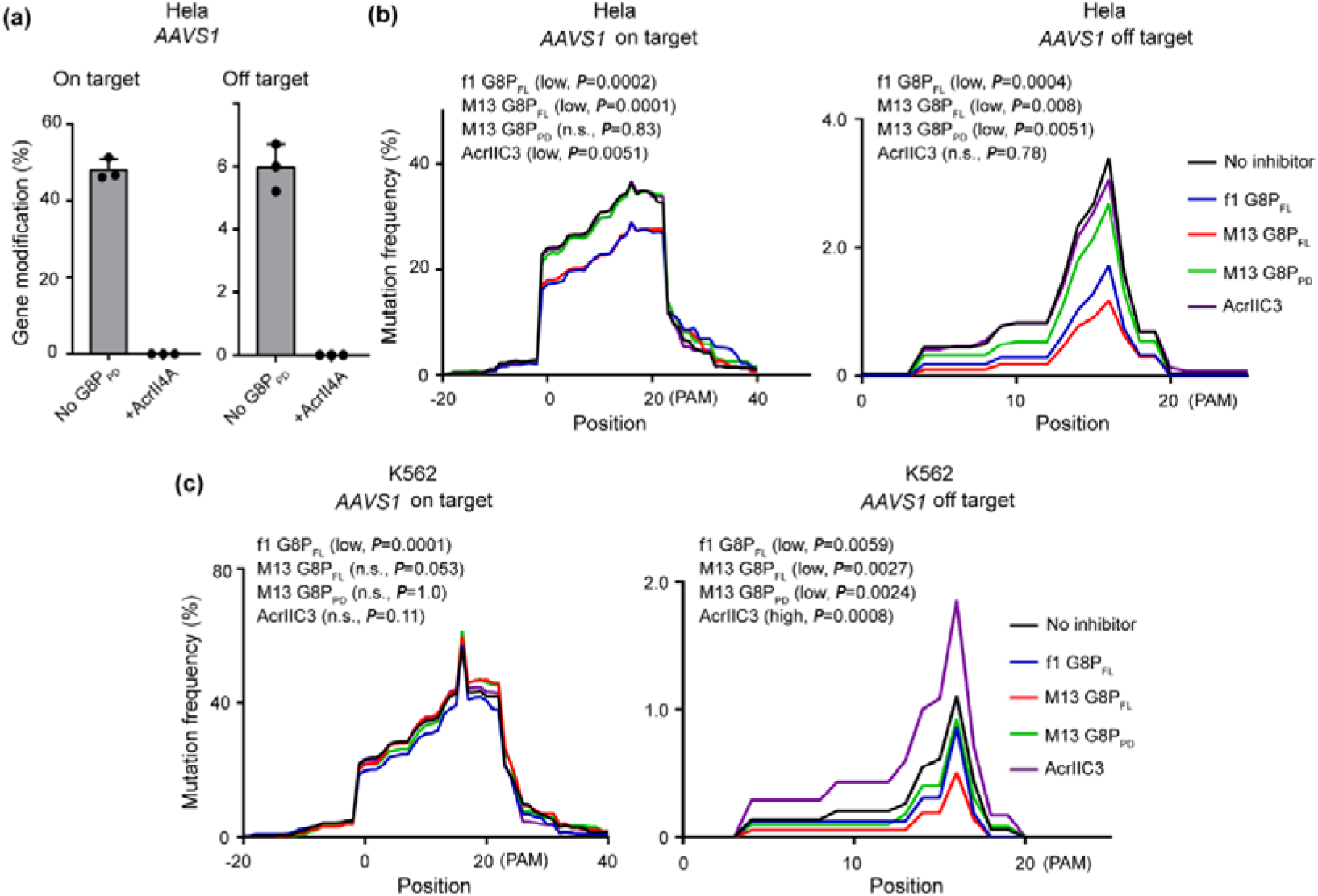
Analyses of the effects of co-transfected G8P and Acrs on the genome-editing activity of SpCas9. (**a**) AcrII4A abolishes the on- and off-activities of SpCas9, as determined by T7E1 assay. The results are shown as mean ± SD (*n* = 3). (**b**-**c**) Density plot showing the distribution of mutation rates along the gene-edited genomic sites of *AAVS1* in Hela (**b**) and K562 (**c**) cells. The mean value of three biological replicates are displayed. Significant difference between test groups and mock is determined by one-way ANOVA with Dunnett’s multiple comparisons test. Low and high indicates increased and decreased mutation rates, respectively. The adjusted *P* values are indicated.

Similarly, M13 G8P_PD_ reduced the off-target events of SpCas9 in K562 cells but retained the on-target activity (Fig. 6c). Co-transfection of M13 G8P_FL_ or f1 G8P_FL_ in K562 cells led to minimum or no significant decrease of the on-target activity of SpCas9 but markedly reduced the off-target events (Fig. 6c). Surprisingly, in K562 cells AcrIIC3 had little effect on the on-target activity of SpCas9 but notably increase the off-target activity (Fig. 6c).

Examination of the effects of co-transfected G8Ps on SpCas9 at another therapeutically relevant genomic site *HBB* [58] illuminated that G8Ps could significantly reduce off-target activity with little perturbation of the on-target activity (Additional file1: Figure S7). Moreover, G8P treatment did not alter the patterns of CRISPR-induced mutations, as characterized the positions of peak mutations (Additional file1: Figure S7). Collectively, these data are in line with the observation at *AAVS1* site and suggest that co-transfection of G8P-based anti-CRISPR agents can reduce the off-target events of constitutively expressed SpCas9 in human cells with minimum perturbation on the on-target activity.

CCK-8 assay revealed little or no change of the cell viability of HEK293 cells that were pre-incubated or co-transfected with CRISPR inhibitors (Additional file1: Figure S8), ruling out the possibility that the observed inhibitory effects of G8Ps are the results of associated cytotoxicity. To explore the possible mechanism of G8P-mediated improvement of CRISPR-Cas9 targeting, we performed chromatin immunoprecipitation-quantitative PCR (ChIP-qPCR) to analyze the effects of G8P on the binding of a catalytically inactive Cas9 (dCas9) at the *AAVS1* on target and pre-determined off-target sites. It was found that the presence of M13 G8P_PD_ or M13 G8P_FL_ did not significantly improve the binding specificity at on-target site (Additional file: Figure S9). This suggested that G8P-mediated improvement of CRISPR-Cas9 targeting was unlikely attributed to improved Cas9 binding. Further studies are required to illustrate the detailed mechanism of action of G8P in living cells.

## Discussion

In this study, we reported the discovery of anti-CRISPR peptides from inoviridae bacteriophages. The clue leading to this discovery was based on the observation that intact M13 phage inhibited the DNA cleavage activity of SpCas9 in *in vitro* reactions. Although we focused the investigation on the major coat protein G8P in the present study, it is likely that other surface-exposed minor coat proteins may attribute to intact phage-mediated SpCas9 inhibition.

Unlike previously described Acrs that inhibit CRISPR-Cas by disrupting DNA binding [36, 45, 59] or DNA cleavage [36], G8P inactivates CRISPR-Cas9 via a distinct mechanism by preventing Cas9 from sgRNA binding. Moreover, to achieve efficient *in vitro* and *in vivo* inhibition, G8P_PD_ must access SpCas9 prior to its binding with sgRNA, indicating that G8P_PD_ binds to apo-Cas9, but not sgRNA-bound Cas9. These results suggest that G8P_PD_ and sgRNA are mutually exclusive for binding with Cas9 nuclease. One straightforward explanation is that G8P_PD_ and sgRNA compete for the same binding pocket. However, MS and mutational studies suggest that the binding site of f1 G8P_PD_ is located on the PI domain of Cas9, distal from sgRNA or DNA binding pockets, thus suggesting against direct competition between G8P_PD_ and sgRNA for the same binding pocket. These results strongly suggest that G8P_PD_ function as allosteric inhibitors to SpCas9.

As allosteric inhibitors, G8P_PD_ may prevent Cas9-sgRNA binding by introducing steric hindrance or conformational changes. It is known that the binding of guide RNA can induce conformational rearrangements of Cas9 [60], thus it is possible that G8P_PD_ or sgRNA binding can transform SpCas9 from a flexible conformation to a closed conformation, which prevents SpCas9 from binding with the other counterpart. Additional experiments are required to better illustrate the mechanism of SpCas9 inhibition by G8P. Collectively, G8P-mediated allosteric inhibition of Cas9 and sgRNA binding represents a unique CRISPR-inactivating mechanism that may have important implications for developing next-generation anti-CRISPR agents.

Existing Acr proteins can display nanomolar binding affinity to Cas9 [36]. By contrast, inoviridae phage G8P_PD_ peptides exhibit weak affinity as evidenced by the micromolar IC_50_ in the *in vitro* cleavage reaction (Fig. 1) and by its inability to completely block the assembly of Cas9 and sgRNA (Fig. 2). Interestingly, intact M13 phage inhibits Cas9 with an IC_50_ of approximately 5 nM, 1000-fold lower than that with G8P_PD_. The increased potency with intact phage may result from the enhanced cooperativity and avidity afforded by the multimeric assembly of the phage capsid that carries 2700 copies of G8P.

Although we have demonstrated that inoviridae phage G8P_PD_ can function as anti-CRISPR peptides, the biological relevance of our discovery is yet to be explored. Under native context, the major coat protein G8P does not enter bacterial cytoplasm during phage infection. It is thus unlikely for G8P to exert anti-CRISPR function at the early stage of phage infection. Instead, G8P could interact with CRISPR-Cas in bacterial cytoplasm after the phage genome is translated. This post-translational inhibitory activity will require G8P-coding DNA to evade CRISPR attack during phage infection. In addition, the SpCas9-inactivating G8P_PD_ discovered in the present study are encoded by phages that infect *Escherichia coli* and *Pseudomonas aeruginosa*, which do not harbor type II CRISPR-Cas system. Horizontal gene transfer could explain for the cross-species CRISPR inactivation [26, 32], however a systemic phylogenetic analysis is necessary to reveal the evolutionary implications of the anti-CRISPR activity of G8Ps.

Despite the elusive biological relevance of the anti-CRISPR activity of G8P, we nevertheless demonstrated that the genome-editing activity of SpCas9 in human cells can be modulated by G8P_PD_. To the best of our knowledge, our discovery represents the first peptides known to exhibit Cas9-inhibiting activity and expands the anti-CRISPR agent toolbox which is currently composed of anti-CRISPR proteins [21], small-molecules [20] and synthetic oligonucleotides [61]. Compared with Acr proteins that are typically 10 to 20 kD in size, G8P_PD_ peptides are small in size and can be chemically synthesized in large scale. The facile manufacturing process is critical for the rapid evaluation of the structure-activity relationship to identify enhanced anti-CRISPR peptides. The varied Cas9-inhibiting activities observed among different inoviridae phage G8P_PD_ supports the notion of improving SpCas9-inhibiting peptides by sequence optimization. In addition, the differential effects of G8Ps on SpCas9, NmCas9 and SaCas9 indicate that it may be feasible to design specific peptide inhibitors for different Cas9 homologs.

G8P_PD_ does not alter the pattern of SpCas9-induced mutations, suggesting that G8P_PD_-mediated Cas9 inactivation does not interfere with the downstream DNA repair pathway. This feature facilitates the therapeutic applications of G8P_PD_ as CRISPR-Cas off-switches by restricting the alterations of genome-editing outcome. It has been demonstrated that increasing the intracellular concentrations of Cas9 can yield elevated off-target activity without further increasing the saturated on-target cleavage, thereby resulting in decreased specificity of genome targeting [12]. It has been proposed that the specificity of SpCas9 in human cells may be increased by partial inhibition with weak CRISPR-Cas inhibitors [20]. In the current study, we found that transfection of strong CRISPR-Cas inhibitor AcrII4A inhibited SpCas9 activity to near complete in human cells when transfected prior to (Fig. 5b) or simultaneously with (Fig. 6a) sgRNA- and SpCas9-coding plasmids. By contrast, co-transfection of G8P with CRISPR-Cas9 showed little inhibition of the on-target activity but reduced the off-target events at endogenous genomic sites (Fig. 6 and Additional file: Figure S7).

Therefore, we propose a two-mode mechanism of action for G8P to suppress SpCas9 activities in living cells: pre-incubation of G8P can saturate the intracellular concentrations of G8P and efficiently inhibit the activity of subsequently transfected Cas9 by competing with sgRNA for binding with apo-Cas9, while co-transfection of G8P with Cas9 and sgRNA will not affect the assembly of Cas9-sgRNA for cleavage at on-target sites but only prevent excess Cas9 from off-target editing (Figure 7). Interestingly, it was observed that the AcrIIC3, a weak inhibitor to SpCas9 (Fig. 5b), did not increase the specificity of SpCas9 in human cells (Fig. 6b-c). This result along with the results of G8P_PD_ suggest that the mechanism of action of the inhibitors may be also important for their effects on the specificity of CRISPR-Cas9. In addition, It has to be noted, however, that the applicability of G8Ps as agents to improve the specificity of CRISPR-Cas9 requires further investigation on additional cell types, genomic loci and genome-wide mutation profile and improved understanding of the mechanism of action of G8P.

**Fig. 7.**
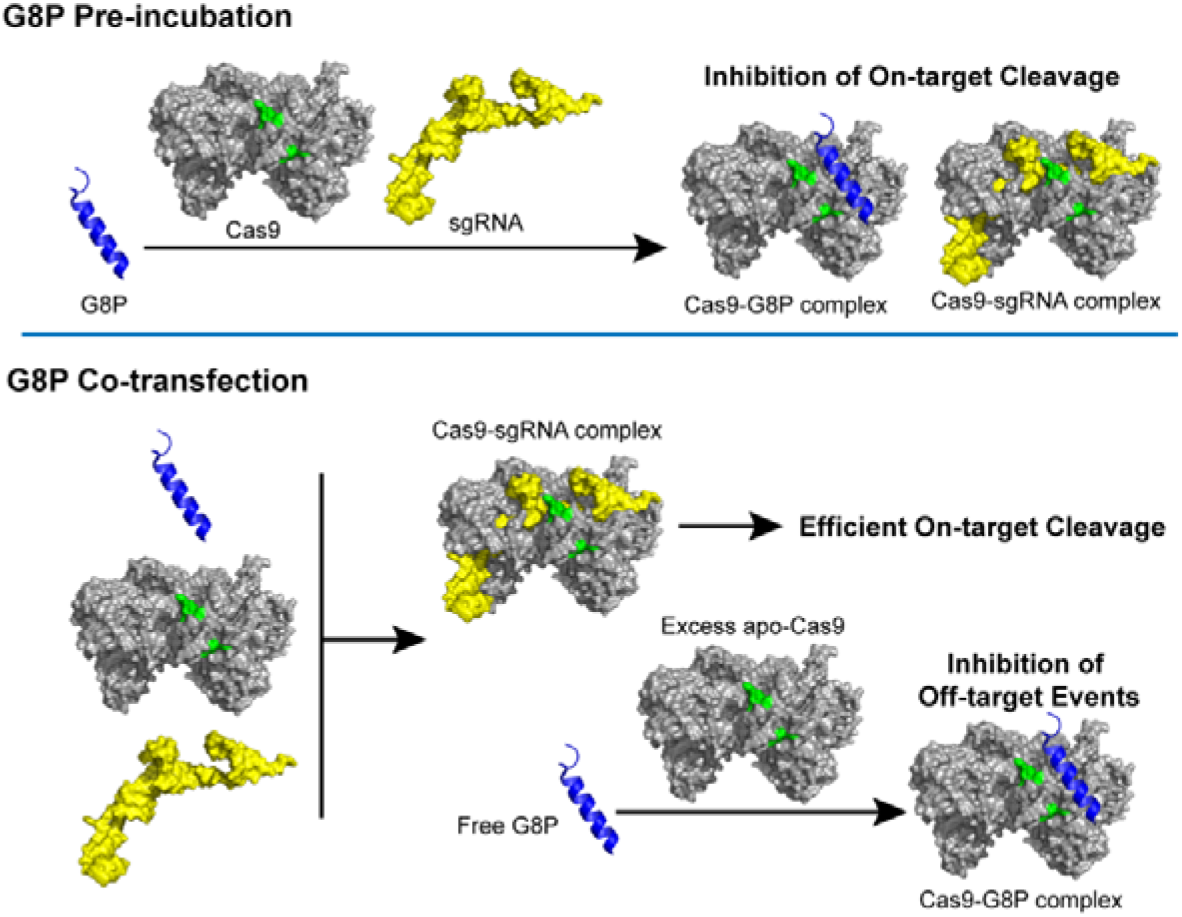
Proposed mechanism of action of G8P in living cells when delivered prior to or simultaneously with Cas9 and sgRNA-coding genes.

## Conclusion

In the present study, we report the surprising discovery of phage-derived peptides that can inhibit the *in vitro* and *in vivo* activities of SpCas9. These peptides inhibited Cas9 activity by disrupting Cas9 and sgRNA binding in an allosteric manner. We show that the genome-editing activity of CRISPR-Cas9 can be harnessed by G8P. This unique mechanism of action of G8P may provide insights into developing anti-CRISPR agents towards genome- or base-editing systems.

## Methods

### Production of M13 bacteriophage

To produce M13 phage, XL1-Blue *E. coli* was inoculated in SB medium supplemented with 2% glucose and grown until OD_600_ reached 0.5. M13 phages were transfected and incubated with XL1-Blue *E. coli* at 37 □ for 30 min and the culture was shaken at 37 □ for 1 h. Thereafter, bacteria were centrifuged twice at 3,000 rpm for 15 min at 4 □ and washed with SB medium to remove glucose and free phage particles. The cells were re-suspended with SB medium and diluted to an OD_600_ of 0.8 in 200 mL culture and grown at 30 □ overnight with shaking. The next day, the culture was centrifuged at 5,000 rpm for 15 min at 4 □ and the supernatant containing phage particles was collected. The supernatant was gently mixed with NaCl-PEG buffer containing 20% PEG and 2.5 M NaCl and kept on ice for 1 h. The PEG-phage supernatant was centrifuged at 9,000 rpm for 30 min at 4 □ The supernatant was discarded and phage pellet was centrifuged again at 9,000 rpm for 5 min at 4 □ to remove residual medium. The precipitated phage particle was re-suspended in phosphate buffered saline (PBS) for further studies.

### Cell culture

HEK293T cells were maintained in Dulbecco’s Modified Eagle’s medium (DMEM, Gibco/Thermo Fisher Scientific, Shanghai, China) supplemented with 10% fetal bovine serum (FBS, Gibco/Thermo Fisher Scientific) at 5% CO_2_ and 37 °C in a fully humidified incubator and were passaged when 70-90% confluency was reached. K562 cells were cultured in RPMI-1640 medium supplemented with 10% FBS, 100 IU/mL of penicillin and 100 μg/mL of streptomycin at 37 °C under 5% CO_2_. Cell lines were validated by VivaCell Biosciences (Shanghai, China). Cell viability was determined using CCK-8 assay ().

### Expression and purification of SpCas9 proteins

pET28b plasmids coding SpCas9 WT, K1158 mutant and K1176 mutant proteins were transformed into *E. coli* BL21 (DE3) cells. Single colonies were picked and grown in 2 liter LB media supplemented with 50 μg/mL kanamycin. Culture was grown to an OD_600_ of 0.8. Protein expression was induced with 0.2 mM isopropyl-β-D-thiogalactopyranoside (IPTG) at 16 °C overnight. Cells from 2 liter culture were pelleted by centrifugation at 6,000 g at 4 °C for 15 min and then re-suspended in 40 mL binding buffer containing 20 mM TrisHCl, pH 8.0 and 0.5 M NaCl. Cell suspension was then supplemented with 1 mM Tris (2-carboxyethyl) phosphine (TCEP) and 1× complete inhibitor cocktail (Roche). Cells were lysed by sonication on ice and then centrifuged at 80,000 g at 4 °C for 30 min. The supernatant of cell lysate was incubated with 1 mL Ni-NTA agarose beads (QIAGEN) at 4 °C for 1 h. The resin was washed with 20 mL wash buffer that was made by supplementing binding buffer with 30 mM imidazole. Proteins were eluted with 5 mL elute buffer that was made by supplementing binding buffer with 300 mM imidazole. Eluted protein samples were further purified by gel filtration using Superose 6 10/300 column (GE Healthcare). These proteins were buffer-exchanged to storage buffer containing 20 mM HEPES, pH 8.0 and 200 mM NaCl, aliquoted and stored at −80 °C.

### Construction of G8P_PD_ overexpression plasmids

Human codon-optimized DNA sequences encoding M13, f1, f1 mutant 2 and pf1 G8P_PD_ were cloned into the BamHI/XbarI sites of pcDNA3.1(+) by plasmid recombination kit Clone Express (Vazyme). These G8P_PD_ peptides carry an N-terminal SV40 nuclear localization signal (NLS) for co-localization with Cas9 proteins. G8P_PD_ peptides were cloned into plv-EF1α-mCherry plasmid harboring mCherry fluorescent protein marker. sgRNA was cloned into pGL3-U6-gRNA plasmid carrying green fluorescent protein (GFP).

### *In vitro* transcription of sgRNA

*CCR5*-targeting sgRNA (Additional file 2: Table S1) was transcribed from a sgRNA-coding PCR product with a 5′ T7 promoter sequence using HiScibe T7 Quick High yield RNA Synthesis kit (NEB). The transcription was performed at 37 °C overnight and then purified by phenol: chloroform extraction, followed by ethanol precipitation. Purified sgRNA was quantified by spectrometry and stored at −80 °C.

### *In vitro* cleavage assay

Cas9 protein and transcribed sgRNA were incubated for 10 min at room temperature in reaction buffer containing 1× NEB buffer 3.1 (NEB Biolabs) supplemented with 1 mM DTT) to form Cas9-sgRNA RNP complex. Cleavage was performed in 10 μL reactions containing 100 ng of substrate DNA and 1 μL RNP complex of indicated concentrations at room temperature for 1 h. Reactions were terminated by addition of 1× DNA loading buffer and resolved on 2% agarose gels. For inhibition experiments, G8P_PD_ peptides (GeneScript, Nanjing, China) were dissolved in deionized distilled water and incubated with Cas9 protein or pre-assembled Cas9-sgRNA RNP for 10 min at room temperature and the mixed solution was then added to the *in vitro* cleavage reaction. For experiments comparing the inhibitory activities of G8P_PD_ peptides, 0.1% dimethyl sulfoxide (DMSO) was included in the reaction solution to solubilize lyophilized peptide samples. His-tagged full-length G8P had poor expression in BL21 (DE3) *E. coli* when expressed from pET28a vector and was thus excluded from the *in vitro* analyses.

### Electrophoresis mobility shift assay (EMSA)

sgRNA concentration is fixed to 15 μM and Cas9 protein was titrated with a molar ratio of Cas9 over sgRNA ranging from 0.05 to 1. f1 G8P_PD_ of 150, 300 and 600 μM was incubated with Cas9 protein or pre-assembled Cas9-sgRNA RNP complex for 20 min at 25 °C and quenched by addition of 1× native DNA loading buffer containing 40 mM Tris, pH 8.2, 40 mM acetate, 1 mM ethylenediaminetetraacetic acid (EDTA), 12.5% (v/v) glycerol, 0.025% (m/v) bromophenol blue. The samples were run on 2% agarose gels. For the order-of-addition experiment, Cas9 and sgRNA concentrations are fixed to 7.5 and 15 μM respectively. SpCas9 was pre-incubated with sgRNA or G8P_PD_ for 20 min, followed by incubation with the counterpart for 20 min at 25 °C.

### Chemical crosslinking and mass spectrometry

Cas9 protein and M13 G8P_PD_ were crosslinked using collision-induced dissociation (CID)-cleavable cross-linker-disuccinimido sulfoxide (DSSO) following the described procedure [55] with minor modification. Cas9 protein and peptides were mixed in PBS and incubated for 30 min at room temperature. Crosslinking was performed for 30 min by adding DSSO (Thermo Scientific) to protein/peptide solution with 1,000 molar excess. The crosslinking reaction was quenched by excess Tris and the crosslinked products were digested with Chymotrypsin. The LC MSn data of digested peptides were collected on Orbitrap Fusion Tribrid (Thermo Scientific) with an on-line NanoLC system and analyzed using CID-MS^2^-MS^3^ strategy as previously described [62]. Monoisotopic mass of parent ions and corresponding fragment ions, parent ion charge states and ion intensities from LC MS^2^ and LC MS^3^ spectra were extracted using Xcalibur v 3.0 (Thermo Scientific). Database searching was performed using Proteome Discoverer v 2.2 software (Thermo Scientific). Chymotrypsin was set as the enzyme with two missed cleavages being allowed as the maximum values. Protein N-terminal acetylation, methionine oxidation (15.995 Da), carbamidomethyl cysteine (57.021 Da), hydrolyzed lysine DSSO (176.014 Da) and lysine DSSO TRIS (279.078 Da) were selected as variable modifications. In addition, to account for the residual crosslinker three defined modifications on uncleaved lysines were chosen including alkene (C_3_H_2_O, 54 Da), sulfenic acid (C_3_H_4_O_2_S,104 Da) and thiol (C_3_H_2_SO, 86 Da) modifications. A false discovery rate (FDR) of 1% was employed to filter out false positive results. The MS, MS^2^ and MS^3^ mass tolerances were set as 10 ppm, 20 ppm and 0.6 Da respectively.

The XlinkX detect program (Thermo Scientific) was used to search MS^2^ data and identify the list of putative DSSO-interlinked products based on their unique DSSO fragmentation patterns. Monoisotopic masses and charges of parent ions measured in MS^3^ scans for those putative cross-linked peptides were further validated and scored by XlinkX. The final results were confirmed by manual inspection of the MS^2^ and MS^3^ spectra, respectively.

### Circular dichroism

Circular dichroism spectroscopy (Chirascan-plus CD spectrometer, Applied Photophysics) was used to probe the peptide conformational changes. 100 μM WT and mutant 2 G8P_PD_ peptides were dissolved in deionized water. The CD data were recorded at 25 °C and the average value of three biological replicates was presented.

### Inhibition of CRISPR-Cas9 activity in human cells by G8P_PD_ overexpression

K562 cells (2 × 10^5^) were harvested, washed with PBS and re-suspended in 20 μL of SF nucleofection buffer (Lonza). G8P_PD_-coding plasmid (1 μg) was mixed with re-suspended K562 cells and nucleofected by Lonza 4D nucleofector with program FF-120. Immediately following the nucleofection, 100 μL pre-warmed RPMI-1640 medium was added into nucleofection cuvettes and the cells were transferred to culture dishes. At 24 post nucleofection, plasmids encoding SpCas9 (500 ng) and *AAVS1* or *EMX1*-targeting sgRNA (250 ng) (Additional file 2: Table S1) were transfected into G8P_PD_-expressing K562 cells by nucleofection as described above.

Low-passage HEK293T cells were seeded into 24-well plates at a density of 150,000 cells per well. The next day, G8P plasmid (1 μg) were transfected into cells using Lipofectamine 3000 (Invitrogen). At 24 h post G8P transfection, plasmids encoding SpCas9 (0.5 μg) and *AAVS1*-targeting sgRNA (250 ng) were co-transfected into G8P-expressing HEK293T cells using lipofectamine.

For G8P pre-incubation experiments, sgRNA-expressing and G8P-expressing plasmids contain green fluorescent protein (GFP) and mCherry reporters, respectively, the expression of which are under the control of promoters independent of sgRNA or G8P expression. For the mock groups without G8P expression, an empty mCherry-expressing plasmid was transfected to control for cell stress. At least 2,000 mCherry and GFP dual positive cells were collected using a BD FACSAria III flow cytometer (BD Biosciences) for subsequent analyses. For co-transfection experiments, K562 and Hela cells were nucleofected with 1 μg of G8P plasmid, 0.5 μg of SpCas9 plasmid and 0.5 μg sgRNA plasmid. At 48 h after transfection, unsorted cells were collected for subsequent analyses.

For NmCas9 and SaCas9 experiments, low-passage HEK293T cells were seeded into 48-well plates at a density of 100,000 cells per well. The next day, f1 G8P_PD_ plasmid (1.5 μg) were transfected into cells using Lipofectamine 3000 (Invitrogen). At 24 h post G8P_PD_ transfection, plasmids encoding SaCas9 (0.75 μg) and *EMX1*-targeting sgRNA or *FANCY*-targeting sgRNA (0.375 μg) were co-transfected into G8P_PD_-expressing HEK293T cells. For NmeCas9, plasmids encoding NmeCas9 (0.75 μg) and *LINC01588*-targeting sgRNA (0.375 μg) were co-transfected into G8P_PD_-expressing HEK293T cells. At 48 h after transfection of SaCas9 or NmeCas9 and sgRNA plasmids, mCherry and GFP dual positive cells were collected as described above for subsequent analyses.

The genomic DNA of collected cells was extracted using QuickExtract DNA Extraction Solution (Epicentre). Genomic PCR reaction was performed using 100 ng genomic DNA, corresponding primers (Additional file 2: Table S2), Phanta Max Super-fidelity DNA Polymerase (Vazyme) or KOD plus (Takara) using a touchdown cycling protocol (30 cycles of 98 °C for 10 s, 68-58 °C for 15 s and 68 °C for 60 s). The PCR products were digested by T7E1 enzyme (NEB), resolved on 2% agarose gel and then analyzed by densitometry measurements as described [63]. Two or three biological replicates were performed for each condition.

The sgRNA sequences used in this study are as follows (PAM sequence uppercased): SpCas9-*CCR5* (tgacatcaattattatacatCGG; for *in vitro* cleavage reaction), SpCas9-*EMX1* (gagtccgagcagaagaagaaGGG), SpCas9-*AAVS1* (gggagggagagcttggcaggGGG), SpCas9-HBB (cttgccccacagggcagtaaCGG), NmCas9-*LINC01588* (cgcaaagctgcatccaccccccgAGACC), SaCas9-*EMX1* (gcaaccacaaacccacgagggCAGAGT), SaCas9-*FANCY* (gcaaggcccggcgcacggtggCGGGGT)

### Next-generation sequencing of edited genomic sites

Genomic DNA (100 ng) from *AAVS1*-edited K562 cells was subject to PCR reactions using stubbed primers (Additional file 2: Table S2). PCR products were purified using Gel Extraction Kit (OMEGA). A high-throughput library preparation kit (Hiseq3000 SBS&Cluster kit) was used to generate dual-indexed sequence. Two or three biological replicates were processed by Genergy Biotech (Shanghai, China) or Genewiz (Suzhou, Jiangsu, China) using Illumina HiSeq 3000 platform.

Next generation sequencing library preparations were constructed following the manufacturer’s protocol (VAHTS Universal DNA Library Prep Kit, Illumina). For each sample, more than 50 ng purified PCR fragment was used for direct library preparation. The fragments were treated with End Prep Enzyme Mix for end repairing, 5′ phosphorylation and dA-tailing in one reaction, followed by a T-A ligation to add adaptors to both ends. Size selection of adaptor-ligated DNA was then performed using VAHTSTM DNA Clean Beads. Each sample was then amplified by PCR for 8 cycles using P5 and P7 primers. Both P5 and P7 primers carry sequences that can anneal with flowcell to perform bridge PCR. In addition, P7 primer carries a six-base index allowing for multiplexing. The PCR products were cleaned up using VAHTSTM DNA Clean Beads, validated using an Agilent 2100 Bioanalyzer (Agilent Technologies, Palo Alto, CA, USA) and quantified by Qubit2.0 Fluorometer (Invitrogen, Carlsbad, CA, USA). Then libraries with different indexes were multiplexed and loaded on an Illumina HiSeq instrument according to manufacturer’s instructions (Illumina, San Diego, CA, USA). Sequencing was carried out using a 2 x 150 paired-end (PE) configuration. Image analyses and base calling were conducted by the HiSeq Control Software (HCS) + OLB + GAPipeline-1.6 (Illumina) on the HiSeq instrument. Sequencing reads were obtained in the Fastq format.

Amplicons with less than 6 M read counts were excluded from the analyses. Short reads were aligned to the reference sequence by Bowtie2 [64] with the following parameters: -D 5 -R 3 -N 1 --gbar 1 --rdg 5,1 --rfg 5,1 --dovetail. Aligned reads were sorted by SAMtools [65] and INDEL and SNP calling was performed by mpileup [66] with maximum read depth per sample equal to the total reads mapped. VarScan v2.4 [67] was used for the quality control of SNPs and INDELs in mpileup output with a minimum variant frequency of ≥ 0.001, and a *P* value threshold of ≤ 0.05. With the above settings, the following items were quantified including the proportions of reads with INDELs/SNPs at each position in the 20 bp target region, the proportions of INDEL with different insertion or deletion length, the proportions of INDEL reading frames (3N, 3N+1 and 3N+2) and the proportions of reads harboring variants over the total number of aligned reads.

### ChIP-qPCR analyses of the effects of G8Ps on dCas9 binding

For ChIP experiments, low-passage Hela cells were seeded on to 15-cm plates at a density of 4×10^6^ cells per plate. The next day, a total of 40.5 μg M13 G8P_PD_ or M13 G8P_FL_ plasmid, 20.25 μg 3×Flag-tagged dCas9 plasmid and 20.25 μg sgRNA plasmid were transfected into cells using Lipofectamine 3000 (Invitrogen). Two biological replicates were performed for each condition. ChIP was performed with Simple ChIP Enzymatic Chromatin IP Kit (Cell Signaling Technology) following the manufacturer’s instructions. After 48 □ h, the cells were washed once with PBS, and 1.25 mL of 16% formaldehyde (Life Technologies) was added to 20 mL of serum-free DMEM for cross-linking to a final formaldehyde concentration of 1%. After 10 min of cross-linking at room temperature, the cells were harvested for the ChIP assay. The chromatin was sonicated using Branson digital sonifier for 10 cycles of “on/off” pulse with 30 s interval. Fragmented chromatin was diluted and incubated with 2 μg anti-Flag Chip grade antibody (Sigma, USA) overnight at 4 °C. qPCR with immunoprecipitation-purified DNA as the template was used to analyze the effects of G8Ps on dCas9 binding at the on-target and off-target sites.

### Statistical analyses

Two or three biological replicates were performed for each experimental condition. Significant difference was analyzed using one-way ANOVA with Dunnett’s multiple comparisons test unless otherwise noted.

## Additional files

Additional file 1: Figure S1-S9.

Additional file 2: Table S1-S2.

## Abbreviations

CRISPR: Clustered regularly-interspaced short palindromic repeats
Cas: CRISPR-associated protein
sgRNA: Single guide RNA
Acrs: Anti-CRISPR proteins
dsDNA: Double-stranded DNA
G8P_PD_: Periplasmic domain of the major coat protein G8P
G8P_FL_: Full-length major coat protein G8P
RNP: Ribonucleoprotein
EMSA: Electrophoresis mobility shift assay
MS: Mass spectrometry
CID: Collision-induced dissociation
DSSO: Disuccinimido sulfoxide
PI domain: PAM-interacting domain
CD: Circular dichroism

## Acknowledgements

We thank the Analytical Platform and High-Throughput Screening Platform at Shanghai Institute for Advanced Immunochemical Studies (SIAIS) at ShanghaiTech University for the support of mass spectrometry and flow cytometry experiments.

## Author contributions

P.M. and J.L. conceptualized study. P.M., J.L., Y.-R.C. and S.-J.W. designed the experiments and analyzed data. Y.-R.C. and S.-J.W. performed the *in vitro* and *in vivo* Cas9-inhibiting experiments. J.C. analyzed next-generation sequencing data. J.Li manufactured M13 phage and initial phage selections. P.M. and S.W performed the crosslink experimenting. W.C. and W.Z. performed MS experiment and analyzed the data. B.M. and B.Y. provided purified WT SpCas9 proteins and advice on the purification of other SpCas9 variants. S.W. and Z.Z. helped to express and purify SpCas9 proteins. B.J. and G.Y. provided critical resources. P.M. and J.L. wrote the manuscript. All authors discussed the results and commented on and approved the manuscript.

## Availability of data and materials

The mass spectrometry data have been deposited into ProteomeXchange with the accession number PXD012466. Deep sequencing data have been deposited into NCBI SRA database with the accession number SRP180801 and SRP199555. Additional data that support the findings of this study are available upon request from the lead contact author J. Liu.

## Ethics approval and consent to participate

Not applicable.

## Consent for publication

Not applicable.

