## Supplemental Figures for "Allosteric Inhibition of CRISPR-Cas9 by Bacteriophage-derived Peptides"

**Additional file 1: Figure S1-S5**

**TABLE OF CONTENT**

1. **Figure S1** Comparison of the *in vitro* activity of M13 and f1 phage G8P_PD_.
2. **Figure S2** MS analyses of the interface between SpCas9 and M13 G8P_PD_.
3. **Figure S3** Construction and purification of SpCas9 K1158 and K1176 mutants.
4. **Figure S4** Profile of SpCas9-induced mutations in the absence and presence of G8P_PD_.
5. **Figure S5** G8P_PD_ peptides derived from inoviridae bacteriophages.
6. **Figure S6** NGS analyses of the effects of G8P pre-incubation on NmCas9 and SaCas9 in HEK293 cells.
7. **Figure S7** NGS analyses of the effects of G8Ps on the specificity of SpCas9 at *HBB* site in HEK293 cells.
8. **Figure S8** Cytotoxicity of CRISPR inhibitors.
9. **Figure S9** ChIP-qPCR analyses of the effects of M13 G8Ps on Cas9 binding at *AAVS1* on target and pre-determined off-target sites in Hela cells.

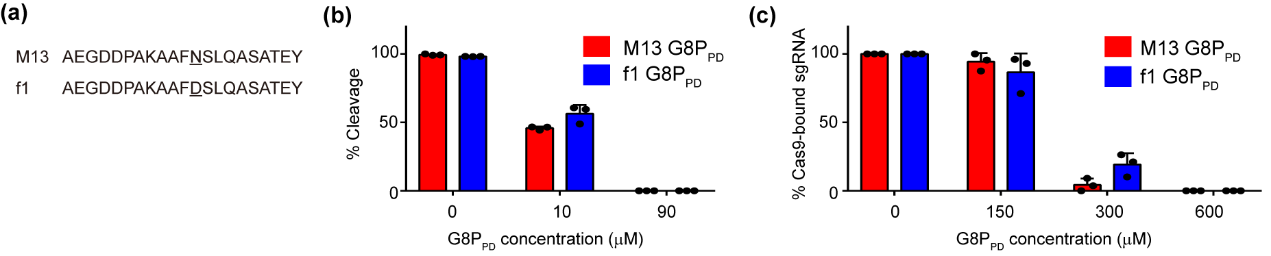

**Figure S1** Comparison of the *in vitro* activity of M13 and f1 phage G8P_PD_. (**a)** Sequence alignment of M13 and f1 G8P_PD_. (**b**) The *in vitro* Cas9-inhibiting activity of M13 and f1 G8P_PD_. M13 and f1 G8P_PD_ of 100 μM were incubated with 50 nM of Cas9 proteins prior to the addition of 50 nM of sgRNA. (c) Inhibition of Cas9-sgRNA assembly by M13 and f1 G8P_PD_ prior to sgRNA addition. Cas9 to sgRNA ratio is fixed to 0.3. The above results (b-c) are shown as mean ± SD (*n* = 3).

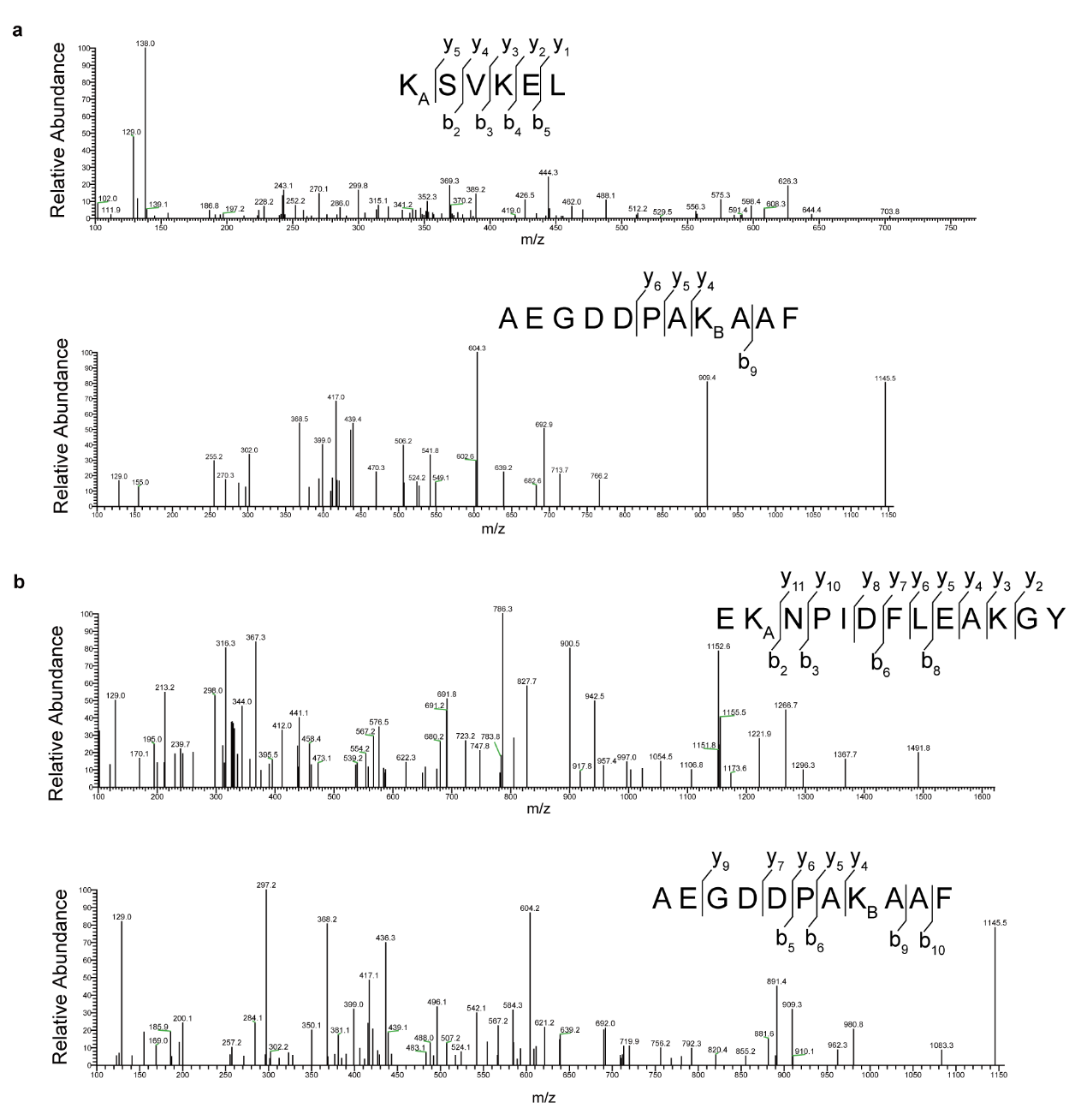

**Figure S2** MS analyses of the interface between SpCas9 and M13 G8P_PD_. (**a**) Tertiary MS for individual peptides KSVKEL and AEGDDPAKAAF as described in Fig. 3b. (**b**) Tertiary MS for individual peptides ENKPIDFLEAKGY and AEGDDPAKAAF as described in Fig. 3c.

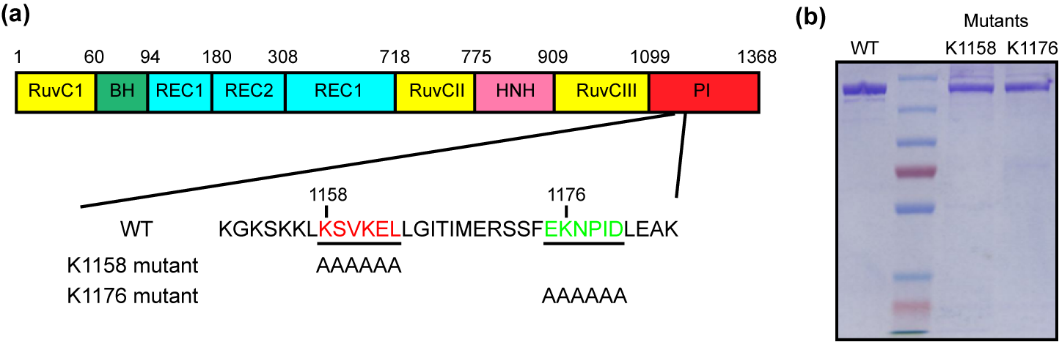

**Figure S3** Construction and purification of SpCas9 K1158 and K1176 mutants. (**a**) Schematic presentation of the structural organization of *S. pyogenes* Cas9. BH, bridging helix. PI, PAM interacting domain. Positions of alanine mutations in K1158 and K1176 mutants are indicated. (**b**) Purified WT, K1158 mutant and K1176 mutant SpCas9 proteins.

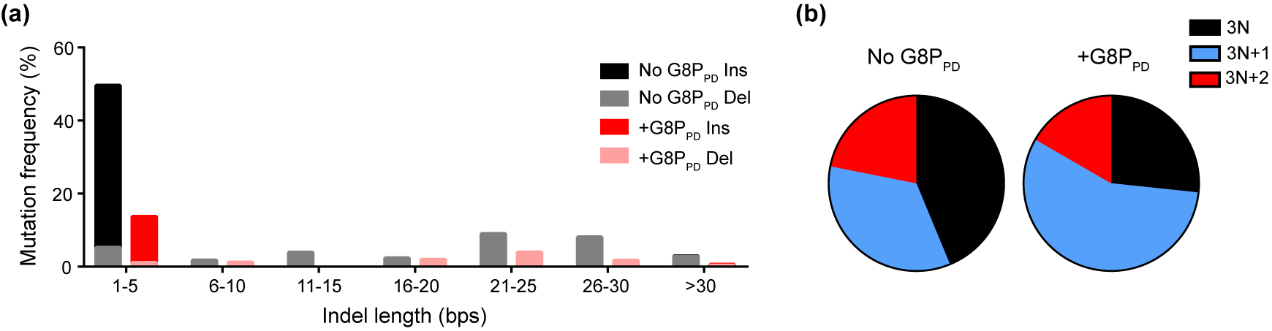

**Figure S4** Profile of SpCas9-induced mutations in the absence and presence of G8P_PD_. (**a**) Distribution of indel length. (**b**) Distribution of indel frame phase calculated as the length of indel modulus. The mean value of two biological replicates are shown.

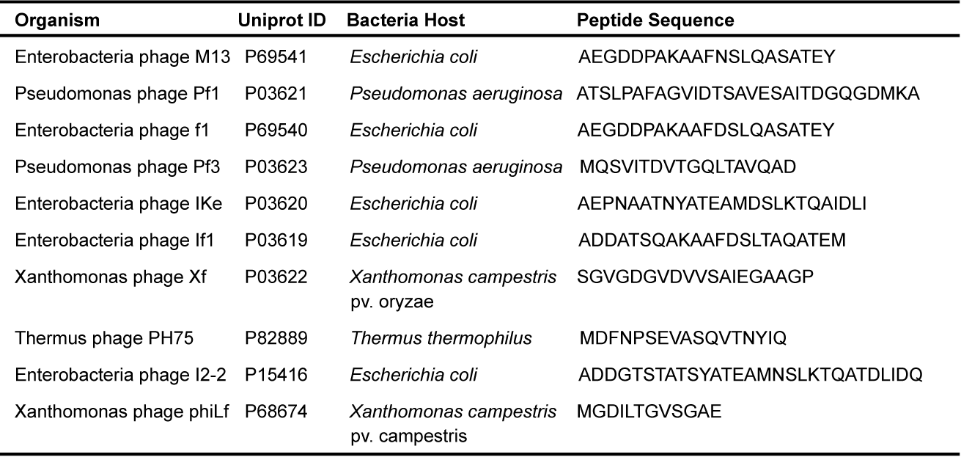

**Figure S5** G8P_PD_ peptides derived from inoviridae bacteriophages.

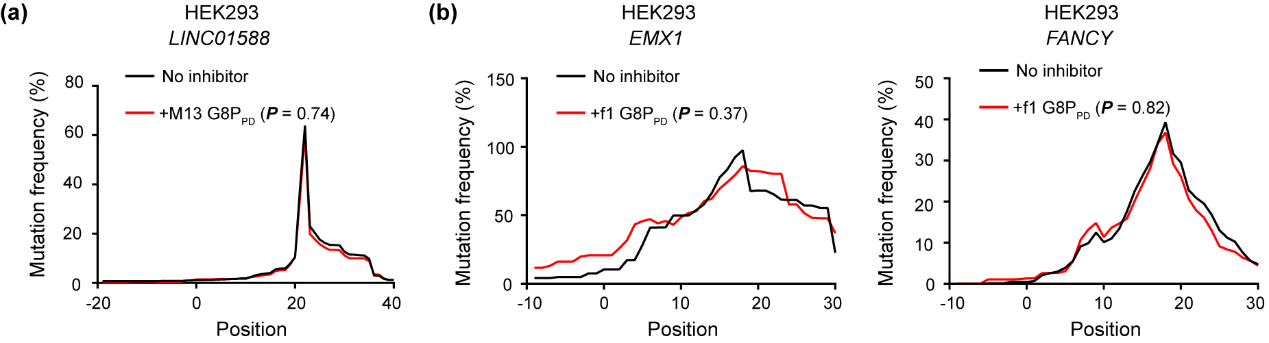

**Figure S6** NGS analyses of the effects of G8P pre-incubation on NmCas9 (a) and SaCas9 (b) in HEK293 cells. The mean value of three biological replicates are displayed. Significant difference between test group and mock is determined by two-tailed, unpaired Student’s *t* test. The adjusted *P* values are indicated.

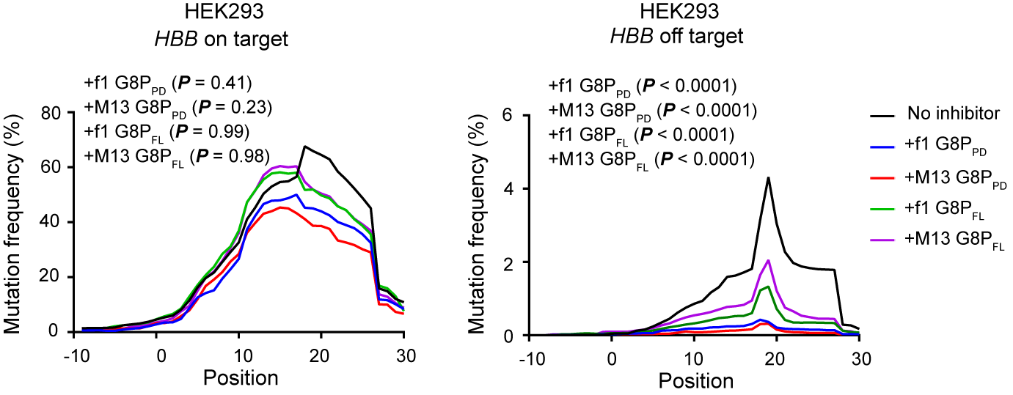

**Figure S7** NGS analyses of the effects of G8Ps on the specificity of SpCas9 at *HBB* site in HEK293 cells. The mean value of three biological replicates are displayed. Significant difference between test groups and mock is determined by one-way ANOVA with Dunnett’s multiple comparisons test. The adjusted *P* values are indicated.

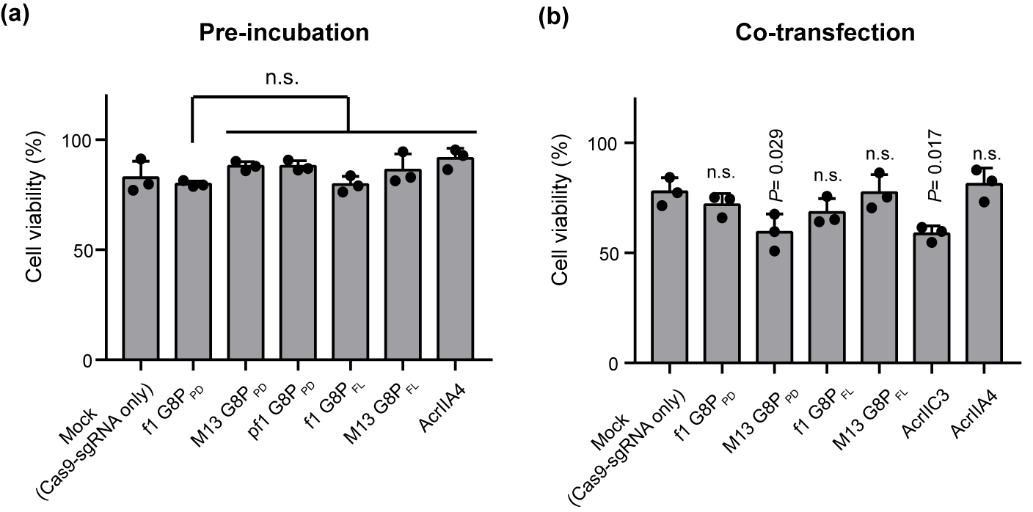

**Figure S8** Cytotoxicity of CRISPR inhibitors. Plasmids encoding CRISPR inhibitors are pre-incubated with or co-transfected in HEK293 cells as described in the Methods in the main text. Cell viability was determined using CCK-8 assay. The results are shown as mean ± SD (*n* = 3). Significant difference between test groups and mock is determined by one-way ANOVA with Dunnett’s multiple comparisons test. The adjusted *P* values are shown. n.s., no significant difference.

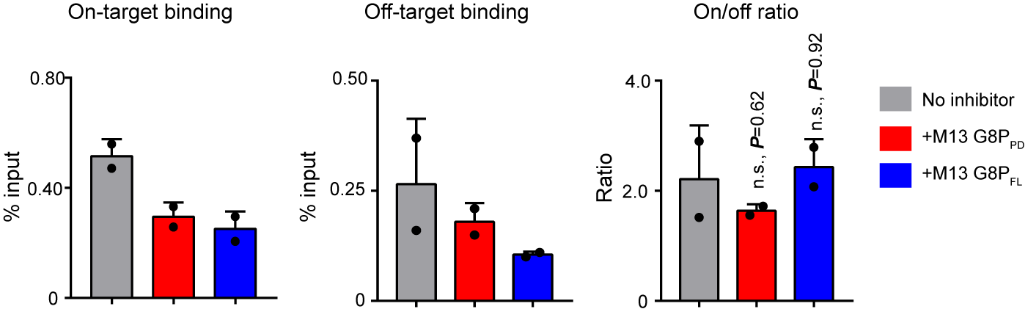

**Figure S9** ChIP-qPCR analyses of the effects of M13 G8Ps on Cas9 binding at *AAVS1* on target and pre-determined off-target sites in Hela cells. The results of two biological replicates are shown as mean ± SD. Significant difference between test groups and mock (no inhibitor) is determined by one-way ANOVA with Dunnett’s multiple comparisons test. The adjusted *P* values are shown. n.s., no significant difference.
