## Supplemental Tables for "Allosteric Inhibition of CRISPR-Cas9 by Bacteriophage-derived Peptides"

**Additional file 2: Table S1-S2**

**TABLE OF CONTENT**

1. **Table S1.** sgRNA-targeted genomic sites in this study.
2. **Table S2.** Primer list.

**Table S1. sgRNA-targeted genomic sites in this study.**

| **Gene** | **Sequence (PAM sequence uppercased)** | **Use** |
| --- | --- | --- |
| SpCas9-*EMX1* | gagtccgagcagaagaagaagggXGG | Genome editing |
| SpCas9-*AAVS1* | gggagggagagcttggcaggXGG | Genome editing |
| SpCas9-*HBB* | cttgccccacagggcagtaaCGG | Genome editing |
| NmCas9-*LINC01588* | cgcaaagctgcatccaccccccgAGACC | Genome editing |
| SaCas9-*EMX1* | gcaaccacaaacccacgagggCAGAGT | Genome editing |
| SaCas9-*FANCY* | gcaaggcccggcgcacggtggCGGGGT | Genome editing |
| SpCas9-*CCR5* | tgacatcaattattatacatCGG | *In vitro* cleavage |

**Table S2. Primer list.**

| **Name** | **Sequence** | **Use** |
| --- | --- | --- |
| CCR5-IVT-FWD | gaaattaatacgactcactataggtgacatcaattattatacatgttttagagctagaaata | *In vitro* transcription |
| CCR5-IVT-REV | agcaccgactcggtgcca | *In vitro* transcription |
| CCR5-Ext-FWD | aggtgagaggattgcttg | Forward external primer for PCR cleavage template |
| CCR5-Ext-REV | aaatgagagctgcaggtg | Reverse external primer for PCR cleavage template |
| CCR5-Int-FWD | gagccaagctctccatctagt | Forward internal primer for PCR cleavage template |
| CCR5-Int-REV | gccctgtcaagagttgacac | Reverse internal primer for PCR cleavage template |
| AAVS1-On-target-NGS-FWD | tctttccctacacgacgctcttccgatctctggtgacacacccccattt | NGS, Forward |
| AAVS1-On-target-NGS-REV | gtgactggagttcagacgtgtgctcttccgatctccaggatcagtgaaacgcac | NGS, Reverse |
| AAVS1-Off-target-NGS-FWD | gcaccttgcaagagaggtac | NGS, Forward |
| AAVS1-Off-target NGS-REV | tgactaaggcagagagaccgaggaagc | NGS, Reverse |
| HBB-On-target-NGS-FWD | gtctccacatgcccagtttc | NGS, Forward |
| HBB-On-target NGS-REV | ctgggcataaaagtcagggc | NGS, Reverse |
| HBB-Off-target-NGS-FWD | taccctttcccgttctccac | NGS, Forward |
| HBB-Off-target NGS-REV | gcacagccagatttgggaat | NGS, Reverse |
| AAVS1-Ext-FWD | ggagttttccacacggacac | Forward external primer for nested PCR for T7E1 assay |
| AAVS1-Ext-REV | cccctatgtccacttcagga | Reverse external primer for nested PCR for T7E1 assay |
| AAVS1-Int-FWD | tgcttctcctcttgggaagt | Forward internal primer for nested PCR for T7E1 assay |
| AAVS1-Int-REV | cggttaatgtggctctggtt | Reverse internal primer for nested PCR for T7E1 assay |
| EMX1-FWD | ggagcagctggtcagagggg | Forward primer for T7E1 assay |
| EMX1-REV | gggaagggggacactgggga | Reverse primer for T7E1 assay |
| SaCas9-EMX1-NGS-FWD | agggctcccatcacatcaac | NGS, Forward |
| SaCas9-EMX1-NGS-REV | ttgtccctctgtcaatggcgg | NGS, Reverse |
| SaCas9-FANCY-NGS-FWD | gatggatgtggcgcaggtag | NGS, Forward |
| SaCas9-FANCY-NGS-REV | aggcgtatcatttcgcggat | NGS, Reverse |
| NmCas9-LINC01588-NGS-FWD | tgcagctatctccgcggccc | NGS, Forward |
| NmCas9-LINC01588-NGS-REV | gtgcggcctgaaatggcaga | NGS, Reverse |
| AAVS1-S3-on-target-ChIP-qPCR-FWD | aacccccacctcctgttagg | ChIP-qPCR, Forward |
| AAVS1-S3-on-target-ChIP-Qpcr-REV | tctctggctccatcgtaagc | ChIP-qPCR, Reverse |
| AAVS1-S3-off-target-ChIP-Qpcr-FWD | aagcgagcattcatgtggaac | ChIP-qPCR, Forward |
| AAVS1-S3-off-target-ChIP-Qpcr-REV | cacccatttgcacaacagcac | ChIP-qPCR, Reverse |
